# Exploring the metabolic profiling of *A. baumannii* for antimicrobial development using genome-scale modelling

**DOI:** 10.1101/2023.09.13.557502

**Authors:** Nantia Leonidou, Yufan Xia, Lea Friedrich, Monika S. Schütz, Andreas Dräger

## Abstract

With the emergence of multidrug-resistant bacteria, the World Health Organization published a catalog of microorganisms urgently needing new antibiotics, with the carbapenem-resistant *Acinetobacter baumannii* designated as “critical”. Such isolates, frequently detected in healthcare settings, pose a global pandemic threat. One way to facilitate a systemic view of bacterial metabolism and allow the development of new therapeutics is to apply constraint-based modelling. Here, we developed a versatile workflow to build high-quality and simulation-ready genome-scale metabolic models. We applied our workflow to create a novel metabolic model for *A. baumannii* and validated its predictive capabilities using experimental nutrient utilization and gene essentiality data. Our analysis showed that our model *i* ACB23LX could recapitulate cellular metabolic phenotypes observed during *in vitro* experiments, while positive biomass production rates were observed and experimentally validated in various growth media. We further defined a minimal set of compounds that increase *A. baumannii* ‘s cellular biomass and identified putative essential genes with no human counterparts, offering novel candidates for future antimicrobial development. Finally, we assembled and curated the first collection of reconstructions for distinct *A. baumannii* strains and analysed their growth characteristics. The presented models are in a standardised and well-curated format, enhancing their usability for multi-strain network reconstruction.

## Introduction

In the 21st century, treating common bacterial infections has become a global health concern. The rapid emergence of pathogens with newly developed resistance mechanisms led to the ineffectiveness of hitherto used antimicrobial drugs. According to their resistance patterns, bacteria are classified into three main categories: multidrug-resistant (MDR, resistant to at least one agent in more than three antibiotic categories), extensively drug-resistant (XDR, non-susceptible to one or two categories), and pandrug-resistant (PDR, non-susceptible to all drugs in all categories)^1^. Pathogens from the last two classes are called “superbugs”. In February 2022, Murray et al. developed predictive statistical models within a large-scale global study and estimated 1.27 million deaths directly associated with antimicrobial resistance (AMR)^2^. The same study underlines the highly virulent ESKAPE pathogens (*Enterococcus faecium, Staphylococcus aureus, Klebsiella pneumoniae, Acinetobacter baumannii, Pseudomonas aeruginosa*, and *Enterobacter* spp.) as the primary cause of AMR-related deaths, while the World Health Organization (WHO) announced in 2017 the urgent need for novel and effective therapeutic strategies against these microorganisms, assigning them the “critical status”.

Over the years, numerous studies highlighted the Gramnegative human pathogen *Acinetobacter baumannii* of substantial concern in hospital environments attributable to its high intrinsic resistance against antimicrobial agents, including biocides^3, 4, 5, 6^. *A. baumannii* (from the Greek word *akínētos*, meaning “unmoved”) is a rod-shaped, non-motile, and strictly aerobic bacterium. It is an opportunistic pathogen whose adaptable genetic apparatus has caused it to become endemic in intensive care units (ICUs), affecting immunocompromised patients, causing pneumonia, bacteremia, endocarditis, and more. Especially the carbapenem-resistant *A. baumannii* poses a serious global threat with high mortality rates^7, 8, 9^. It targets exposed surfaces and mucous tissues, colonises the human nose^10, 11, 12^ and is closely related to Severe Acute Respiratory Syndrome Coronavirus 2 (SARS-CoV-2) infections^13, 14, 15, 16^. The skin has shown to be a community reservoir for *A. baumannii* in a very small percentage of samples^17, 18^, while its prevalence in the soil is a frequent misconception as species from the genus *Acinetobacter* are ubiquitous in nature^5, 19^. Finally, it shows susceptibility to commonly used drugs, like *β* -lactams, aminoglycosides, and polymyxins.

Systems biology, and especially the field of genome-scale metabolic network analysis, is the key to exploring genotypephenotype relationships, better understanding mechanisms of action of such threatening pathogens, and ultimately developing novel therapeutic strategies. Genome-scale metabolic models (GEMs) combined with constraint-based modelling (CBM) provide a well-established, fast, and inexpensive *in silico* framework to systematically assess an organism’s cellular metabolic capabilities under varying conditions having only its annotated genomic sequence^20^. As of today, they have numerous applications in metabolic engineering, leading to the formulation of novel hypotheses driving the detection of new potential pharmacological targets^21^.

It has been more than a decade since the release of the first mathematical simulation of *A. baumannii* metabolism. Kim et al. integrated biological and literature data to manually build AbyMBEL891, representing the strain AYE^22^. This model was further employed as an essential foundation for future reconstructions; however, its non-standardised and missing identifiers limited its use. Following a tremendous increase in the amount of literature and experimental data regarding *A. baumannii* (over 5,670 articles published between 2010 and 2017 according to PubMed), two novel strain-specific metabolic networks arose, *i*LP844^23^ and the AGORA (Assembly of Gut Organisms through Reconstruction and Analysis) model^24^. Both models were reconstructed in a semi-automated process and simulated the metabolism of two distinct strains: ATCC 19606 and AB0057, respectively. With the help of transcriptomic data of sampled colistin responses and *i*LP844, it was observed that the type strain ATCC 19606 underwent metabolic reprogramming, demonstrating a stress condition as a resistance mechanism against colistin exposure. Alterations in gene essentiality phenotypes between treated and untreated conditions enabled the discovery of putative antimicrobial targets and biomarkers. Moreover, the model for AB0057 was part of an extensive resource of GEMs built to elucidate the impact of microbial communities on host metabolism. The amount of massand charge-balanced reactions in these models is very high; however, they carry few to no database references. Norsigian et al. improved and expanded AbyMBEL891 to finally create the high-quality model *i*CN718 that exhibited a prediction accuracy of over 80 % in experimental data^25^, while Zhu et al. built a GEM for ATCC 19606 (*i*ATCC19606) integrating multi-omics data^26^. Compared to *i*LP844, *i*ATCC19606 incorporates metabolomics data together with transcriptomic data enabling the deciphering of bactericidal activity upon polymyxin treatment and the interplay of various metabolic pathways. Last but not least, in 2020, the first *in vivo* study on *A. baumannii* infection was published utilizing constraint-based modelling^27^. This time, the collection of strain-specific models was enriched with the first GEM for the hyper-virulent strain AB5075 (*i*AB5075). The model was validated using various experimental data, while transcriptomics data was leveraged to identify critical fluxes leading to mouse bloodstream infections. Our literature search revealed one last metabolic model of *A. baumannii* ATCC 17978, named *i*JS784, which, by the time of writing, has not been officially published in a scientific journal or been deposited in a mathematical models database. Instead, it is currently available solely in the form of a dissertation^28^. Nonetheless, the model cannot produce biomass even when all uptake reactions are open and all medium nutrients are available to the cell, making it unusable and hampering reproducibility.

We expanded this collection by building a novel GEM for the nosocomial strain ATCC 17978, named *i*ACB23LX. The presented model follows the FAIR data principles and community standards and recapitulates experimentally-derived phenotypes with high predictive capability and accuracy scores. We enriched the model with numerous database cross-references and computationally inferred the minimal nutritional requirements. Moreover, we used this model to investigate the organism’s growth ability in defined media and a medium simulating human nasal secretions while we assessed its ability to predict essential genes using two different optimization approaches. Among the examined strains, ATCC 17978 is one of the most well-studied, with a substantial amount of experimental data available that can be used to direct model refinement and validation. Besides that, we systematically refined and evaluated all pre-existing reconstructions’ performance to finally create the first compendium of curated and standardised models for *A. baumannii*. With this, we aim to promote further studies to give new insights into this pathogen and promote strain- and species-specific therapeutic approaches.

## Results

### Reconstruction process of the novel metabolic network *i*ACB23LX

To build a high-quality model for *A. baumannii* ATCC 17978, we developed a workflow shown in Figure 1 adhering closely to the community standards^29^ (see *Materials and Methods*). We named the newly reconstructed network *i*ACB23LX, where *i* stands for *in silico*, ACB is the organismand strainspecific three-letter code from the Kyoto Encyclopedia of Genes and Genomes (KEGG)^30^ database, 23 the year of reconstruction, and LX the modellers’ initials. Our protocol involves eight major stages starting from the attainment of the annotated genomic sequence until the model validation, applies to any organism from the tree of life (Archaea, Bac-teria, and Eukarya), and ensures the good quality and correctness of the final model. CarveMe^31^ was used to build a preliminary model, which was subsequently extended and curated manually. We resolved syntactical issues and mass and charge imbalances during manual refinement while we defined missing metabolite charges and chemical formulas. Our final model contains no mass-imbalanced reactions and only two charge-imbalanced reactions. After extensive efforts, resolving all charge imbalances was impossible since all participated metabolites are interconnected to multiple reactions within the network, and any modification in their charge resulted in newly introduced imbalances. The model extension process involved incorporating missing metabolic genes considering the network’s connectivity. Dead-end and orphan metabolites do not exist biologically in the species, implying knowledge gaps in metabolic networks. Moreover, reactions including such metabolites are not evaluated in flux balance analysis (FBA). Hence, reactions with zero connectivity and no organism-specific gene evidence were omitted from the gap-filling. We extended the draft model by 138 reactions, 77 genes, and 110 metabolites in three compartments (cytosol, periplasm, and extracellular space). All in all, *i*ACB23LX comprises 2,321 reactions, 1,660 metabolites, and 1,164 genes (Figure 2). It is the most comprehensive model, while its stoichiometric consistency lies at 100 % and contains no unconserved metabolites. Over 1,800 reactions have a gene-protein-reaction associations (GPR) assigned, while 149 are catalysed by enzyme complexes (GPR contains at least two genes connected via a logical AND).

**Figure 1.**
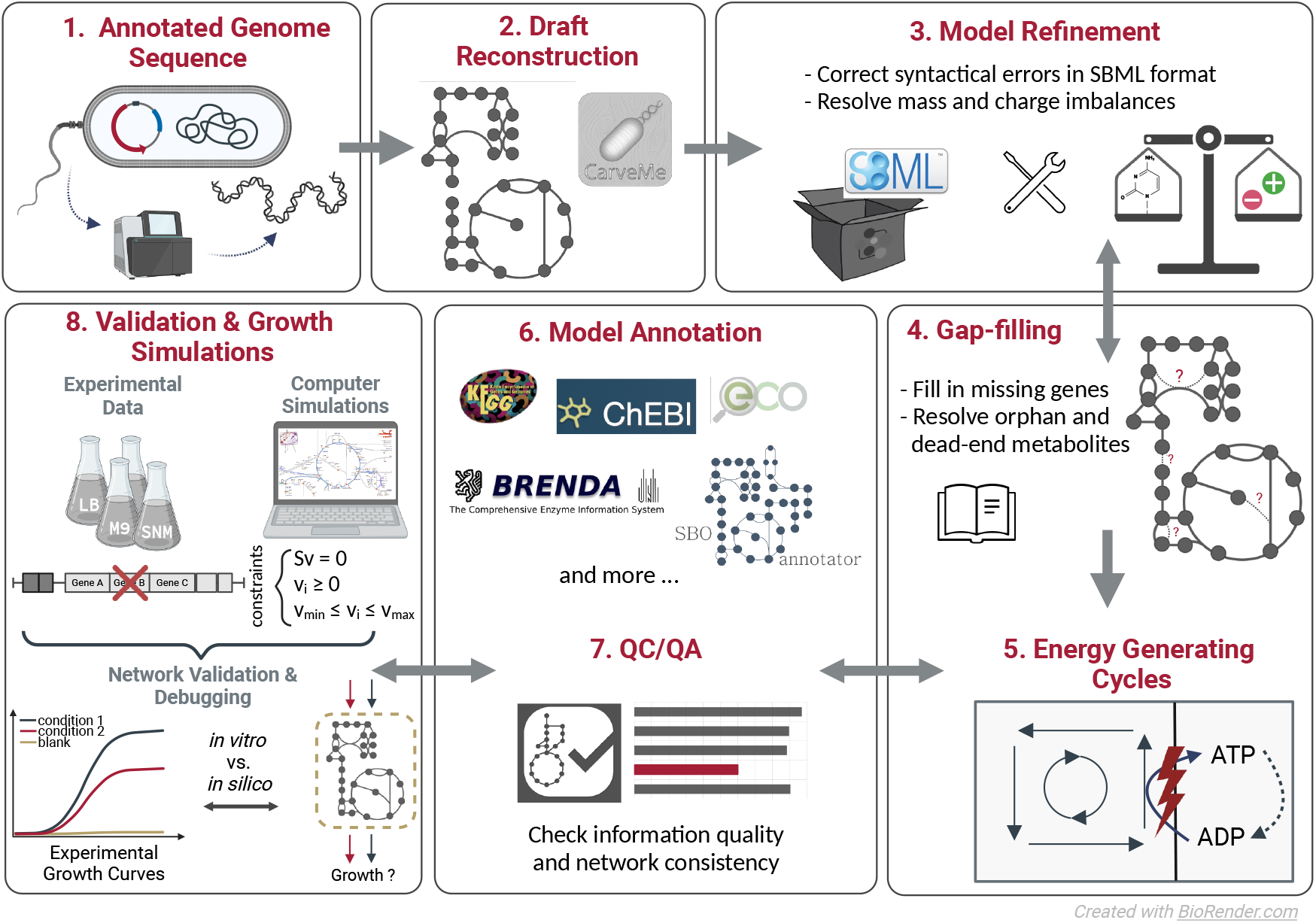
Workflow developed for the metabolic network reconstruction of *i*ACB23LX. The created workflow consists of eight main steps: extraction of the annotated genome, draft model reconstruction, model refinement, gap-filling, investigation of energy-generating cycles, model annotation, quality control and quality assurance (QC/QA), and model validation using experimental data. The growth simulations include the examination of growth requirements and the definition of a minimal growth medium. The last six processes are continuously iterated until the model is of good quality and recapitulates known phenotypes.

**Figure 2.**
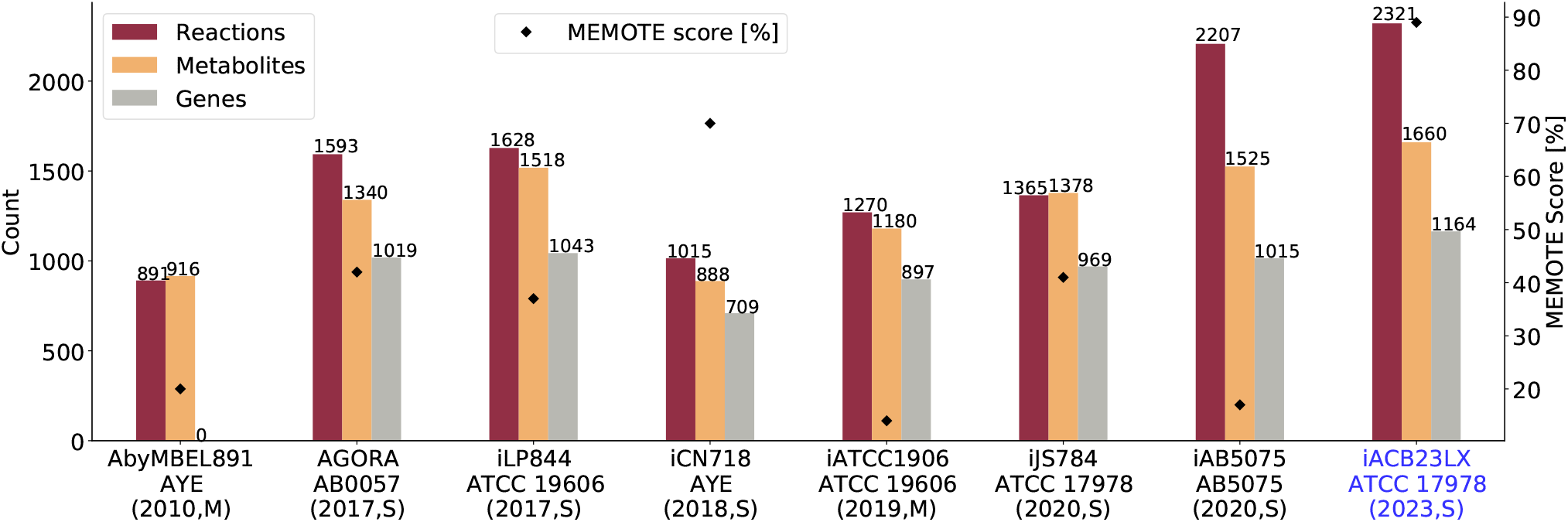
Properties of all metabolic networks of *A. baumannii*. Blue highlights the novel reconstruction for the strain ATCC 17978 presented in this publication. The ordinate represents the MEMOTE scores. The reconstruction process is divided into manual (M, nocomputational tool was used to reconstruct and refine the model) and semi-automated (S, draft obtained via an automated reconstruction tool, while further extension was done manually) and is written together with the publication year. Also, the respective strains annotate the abscissa labels. Our new model exhibits the highest quality score and is more comprehensive and complete than the preceding reconstructions.

Furthermore, we tested our model for energy-generating cycles (EGCs) to prevent having thermodynamically infeasible internal loops that bias the final predictions^32^. We defined energy dissipation reactions (EDRs) for 15 energy metabolites and evaluated their production with FBA after disabling all nutrients from entering the system (see *Materials and Methods*). In our final model, *i*ACB23LX, none of the tested metabolites could be produced; thus, no EGCs are contained. As shown in Figure 1, a plethora of database cross-references was embedded in the model, while Systems Biology Ontology (SBO) terms were defined for every reaction, metabolite, and gene (Figure 3)^34^. Additionally, each reaction was mapped to an Evidence and Conclusion Ontology (ECO) term representing the confidence level and the assertion method.

**Figure 3.**
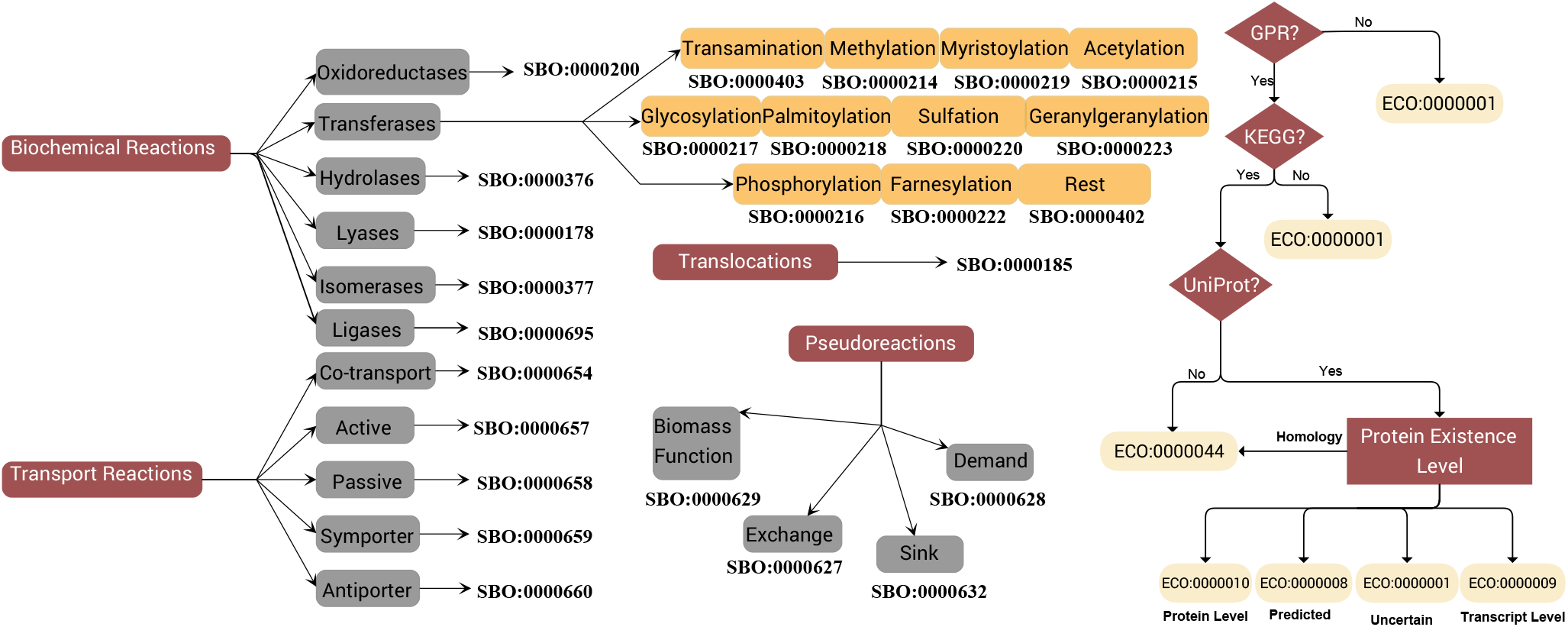
Schematic representation of the SBO and ECO terms mapping. It follows the graphs defined in the repository for biomedical ontologies Ontology Lookup Service (OLS)^33^. The SBO terms were added using the SBOannotator tool^34^. The ECO terms annotated metabolic reactions and were declared based on the presence of GPR along with KEGG and UniProt annotations. Providing UniProt identifiers, the Protein Existence Level guides the mapping to appropriate ECO terms. Figure created with yEd^35^.

To assess the model quality, Metabolic Model Testing (`MEMOTE`)^36^ and the Systems Biology Markup Language (SBML)^37^ Validator from the libSBML^38^ were used. Our metabolic network, *i*ACB23LX, has a `MEMOTE` score of 89 % and all syntactical errors are resolved. Our model undoubtedly exhibits the highest quality score among its predecessors (Figure 2). It must be noted here that the `MEMOTE` testing algorithm considers only the parent nodes of the SBO directed acyclic graph and not their respective children. Assigning more representative SBO terms (Figure 3) does not increase the final score but reduces it by 2 %.

The model is available as a Supplementary File in the latest SBML Level 3 Version 2^39^ and JavaScript Object Notation (JSON) formats with the flux balance constraints (fbc) and groups plugins available.

### *i*ACB23LX is of high quality and exhibits an increased predictive accuracy

#### Prediction and experimental validation of bacterial growth on various nutritional environments

Constraint-based modelling approaches such as FBA estimate flux rates with which metabolites flow through the metabolic network and predict cellular phenotypes for multiple growth scenarios. *A. baumannii* is known to be strictly aerobic and, compared to the majority of *Acinetobacter* species, it is not considered ubiquitous in nature. As a nosocomial pathogen, it has been mostly detected in hospital environments, particularly in the ICUs, and within the human nasal microbiota^10, 11, 12^. We examined various growth conditions to ensure that our model *i*ACB23LX recapitulates these already known and fundamental phenotypes.

First, we tested our model’s capability to simulate a strictly aerobic growth. For this purpose, we examined the directionality of all active oxygen-producing and -consuming reactions when the oxygen uptake was disabled (see Supplementary Figure S1). We observed an accumulation of periplasmic oxygen by reactions that carried remarkably high fluxes and resulted in growth when the oxygen import was turned off. We examined each reaction individually and removed those without gene evidence to correct this. More specifically, we removed the periplasmic catalase (`CATpp`), one of bacteria’s main hydrogen peroxide scavengers. This enzyme is typically active in the cytosol^40^ and was not part of any precursor *A. baumannii* GEM or was found only in cytosol (*i*LP844^23^). To fill the gap and enable the usage of the periplasmic hydrogen peroxide, we included the reaction phenethylamine oxidase (`PEAMNOpp`) in the model. Eventually, our model *i*ACB23LX demonstrated growth only in the presence of oxygen using a rich medium (all exchange reactions are open).

Furthermore, we determined the minimal number of metabolites necessary for growth using *i*ACB23LX and the M9 minimal medium (M9) as a reference. Minimal growth media typically consist of carbon, nitrogen, phosphorus, and sulfur sources, as well multiple inorganic salts and transition metals. These metals are crucial for the growth and survival of all three domains of life; however, they can be transformed into toxic compounds in hyper-availability^41^. The exact composition of our minimal medium (*i*MinMed) is shown in Table 1. It comprises nine transition metals, acetate as the carbon source, ammonium as a nitrogen source, sulfate as a sulfur source, and phosphate as a phosphorus source. Oxygen is also a vital component, as *A. baumannii* is known to be a strictly aerobic bacterium. Previous studies have highlighted the importance of the nutrient metals for *A. baumannii* to survive within the host. More specifically, the bacterium utilises these metals as co-factors for vital cellular processes^42^. Manganese and zinc have also been studied as essential determinants of host defense against *A. baumannii*-acquired pneumonia through their sequestering by calprotectin via a type of bonding called chelation^43^. For the computational simulations, growth rates below 2.81 mmol/(g_DW_ *·*h) were considered to be realistic since the doubling time of the fastest growing organism *Vibrio natriegens* is 14.8 minutes that correspond to 2.81 mmol/(g_DW_*·*h)^36^. Table 2 displays the predicted growth rates of *i*ACB23LX in the respective culture media. In the LB medium, our model grew with the highest rate; 0.5926 mmol/(g_DW_ *·*h). With our self-defined minimal medium, *i*MinMed, our model exhibited the lowest rate; 0.2097 mmol/(g_DW_ *·*h). Notably, our experimental validation revealed similarity between the growth rates obtained from the *in vitro* respiratory curves (Table 2 and Supplementary Figure S2) and those predicted by our *in silico* simulations. Additionally, we examined the growth rate of our model in a rich medium, in which all nutrients are available to the model. With this, the flux through the biomass production was the highest, 2.1858 mmol/(g_DW_*·*h), as expected. This is still less than the growth rate of the fastest organism, increasing the confidence in our model’s consistency and simulation capabilities. Initially, *i*ACB23LX could not predict any realistic growth rate for the simulated media. Using the gap-filling function of CarveMe^31^, we detected three enzymes whose addition into the metabolic network resulted in successful growth in all tested media. These reactions are `PHPYROX`, `OXADC`, and `LCYSTAT`.

**Table 1.** Composition of the computationally defined minimal growth medium *i*MinMed. It consists of nine transition metals, a carbon source, a nitrogen source, a sulfur source, and a phosphorus source. Oxygen was used to simulate aerobic conditions.

|  | Molecular Formula | Name |
| --- | --- | --- |
| Carbon source | C<sub>2</sub>H<sub>3</sub>O<sub>2</sub><sup>-</sup> | Acetate |
| Nitrogen source | NH<sub>4</sub><sup>+</sup> | Ammonium |
| Sulfur source | SO<sub>4</sub><sup>2-</sup> | Sulfate |
| Phosphorus source | HPO<sub>4</sub><sup>2-</sup> | Phosphate |
|  | Ca<sup>2+</sup> | Calcium |
|  | Cl<sup>-</sup> | Chloride |
|  | Cu<sup>2+</sup> | Copper |
|  | Fe<sup>3+</sup> | Ferric iron |
| Transition metals | Co<sup>2+</sup> | Cobalt |
|  | K<sup>+</sup> | Potassium |
|  | Mg<sup>2+</sup> | Magnesium |
|  | Mn<sup>2+</sup> | Manganese |
|  | Zn<sup>2+</sup> | Zinc |
| Oxygen source | O<sub>2</sub> | Oxygen |

**Table 2.** Simulated and empirical growth rates of ATCC 17978 in various growth media. The tested media are the computationallydefined minimal medium (*i*MinMed), the Luria-Bertani (LB) medium, and the synthetic nasal medium (SNM). Growth rates are given in mmol/(gDW*·*h), while doubling times are computed in minutes. The media formulations are available in the Supplementary File S1.

|  | Growth Rates |  | Doubling Times |  |
| --- | --- | --- | --- | --- |
|  | *in silico* | *in vitro* | *in silico* | *in vitro* |
| *iMinMed* | 0.2097 | 0.5402 | 198.32 | 76.70 |
| LB | 0.5926 | 0.7369 | 70.18 | 56.44 |
| SNM | 0.5099 | 0.3592 | 81.56 | 115.78 |

#### *Functional validation of* i*ACB23LX using nutrient utilization data*

Multiple *in silico* approaches have hitherto been employed to predict lethal genes and to assess growth metrics on different carbon/nitrogen sources for severe pathogenic organisms including *Mycobacterium tuberculosis*^44, 45^ and *Staphylococcus aureus*^46, 47^. In 2013 and 2014, two studies were published that examined the phenome and gene essentialities of the strain ATCC 17978^48, 49^. We used these datasets to evaluate the overall performance (functionality and accuracy) of *i*ACB23LX.

Our first validation experiment evaluated the accuracy of our model’s carbon and nitrogen catabolism potentials. More specifically, we detected compounds that could serve as sole carbon and nitrogen sources using the large-scale phenotypic data by Farrugia et al.^48^. Although the authors tested more compounds in total, we could examine only 80 compounds as carbon sources and 48 metabolites as sole nitrogen sources. For the remaining molecules, either no Biochemical, Genetical, and Genomical (BiGG) identifier could be found, or they were not part of the metabolic network. According to the experimental protocol followed by Farrugia et al., we applied the M9 medium and enabled D-xylose as the sole carbon source for the nitrogen testings. As D-xylose was initially not part of the reconstructed network, we conducted extensive literature and database search to include missing reactions. This improved the prediction accuracy, especially for the carbon sources, where an amelioration of 19 % was achieved. In more detail, despite the comprehensive manual curation, the first draft model was reconstructed using the automated tool CarveMe^31^. This resulted in the incorrect inclusion of transport reactions, which were consequently removed to reduce the number of false positive predictions. In both cases, our main objective was to improve the accuracy while keeping the number of orphan and dead-end metabolites low and removing only reactions with no gene evidence (lack of assigned GPR). Similarly, missing reactions were identified and included in the network to eliminate the false negative predictions. For instance, in accordance with the phenotypic data, the strain ATCC 17978 should not be able to grow when utilizing D-trehalose as the sole carbon source. Our model initially predicted a growth phenotype for this carbon source. To overcome this conflict, we deleted the reaction `TREP` with no organism-specific gene evidence, meaning no GPR was assigned. However, it was not feasible to resolve all inconsistencies since adding transport reactions to resolve false positives or false negatives in the nitrogen testings led to more false predictions in the carbon sources. More specifically, when adenosine, inosine, L-homoserine, and uridine are utilised as sole carbon sources, the model should not predict growth, while sole nitrogen sources should result in a nonzero objective value. In this case, adding transporters would resolve false predictions in the nitrogen tests, while it would have induced more false predictions in the carbon sources tests. All in all, *i*ACB23LX exhibited an overall accuracy of 86.3 % for the carbon and 79.2 % for the nitrogen sources test (Figure 4c); however, after further curation, the accuracy was remarkably improved and reached 87.5 %. By adding their corresponding transport reactions, we resolved discrepancies regarding uridine, inosine, adenosine, and L-homoserine.

**Figure 4.**
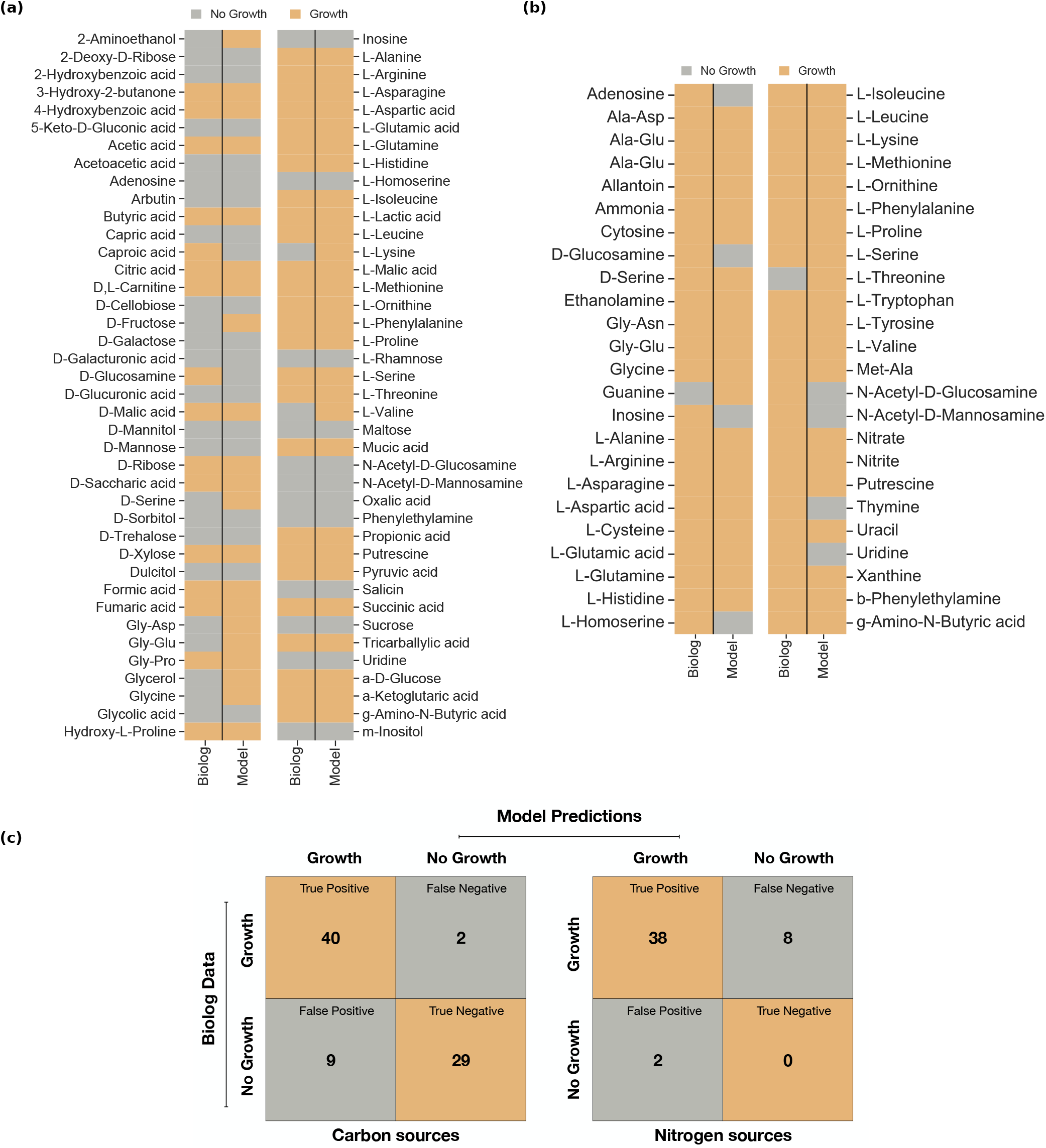
Model predictions compared to the Biolog experimental measurements for various carbon and nitrogen sources. From the Biolog data, only substances mappable to model metabolites were included, while the M9 medium was applied. a) and b) The model’s ability to catabolise various carbon and nitrogen sources was assessed using the strain-specific phenotypic data by Farrugia et al.^48^. Grey indicates no growth, and orange indicates growth. Totally, 80 and 48 compounds were tested as sole carbon and nitrogen sources, respectively. Out of these, 69 and 38 phenotypes were recapitulated successfully by *i*ACB23LX. c) Confusion matrices of model predictions and Biolog experimental measurements. The overall accuracy of *i*ACB23LX is 86.3 % for the carbon (left matrix) and 79.2 % for the nitrogen (right matrix) testings. Orange represents correct predictions, and grey represents wrong predictions.

Our model was able to catabolise 49 sole carbon and 40 sole nitrogen sources (see Figure 4 (a) and Figure 4 (b), recapitulating totally 69 and 38 experimentally-derived phenotypes, respectively.

We further assessed the ability of our model to predict known gene essentialities. First, 1,164 *in silico* single gene deletions were conducted on both LB and rich growth media, respectively, to identify all lethal gene deletions. Subsequently, the ratio between the growth rate after and before the respective knockouts (*FC*_*gr*_) was calculated, and the genes were accordingly classified (see *Materials and Methods*). For the optimization, two mathematics-based approaches from the Constraints-Based Reconstruction and Analysis for Python (COBRApy)^50^ package were deployed: the FBA^51^ and the Minimization of Metabolic Adjustment (MOMA)^52^. Between the two methods, a similar distribution of the *FC*_*gr*_ values was observed (Figure 5 (a) and Figure 5 (b)). Using FBA, 97, 75, and 991 genes were predicted to be essential, partially essential, and inessential on the LB medium, respectively, whereas optimization with MOMA resulted in 110, 85, and 968 genes (Figure 5 (c) and Supplementary Files S2 and S3). These genes were primarily associated with the biosynthesis of cofactors and vitamins, the amino acid/nucleotide metabolism, the energy metabolism, and the metabolism of terpenoids and polyketides. Additionally, we examined in more detail how nutrition availability impacts the gene essentiality by conducting single-gene knockouts in the rich medium. Both optimization methods resulted in more essential genes when the model was required to alter its metabolic behavior due to the absence of nutrients, i.e., with the LB growth medium, compared to the rich medium (Figure 5 (c) and Supplementary Files S2 and S3). In general, FBA detected more genes to be dispensable for growth in both nutritional environments. On the other hand, MOMA classified more genes as essential or partially essential (Figure 5 (c) and Supplementary Files S2 and S3), while genes from FBA build a subset of the essential genes derived by MOMA. Furthermore, we validated the prediction accuracy of *i*ACB23LX using already existing gene essentiality data. At the time of writing, the transposon mutant library by Wang et al. is the only ATCC 17978-specific experimental dataset^49^. With this dataset together and the LB medium, our model demonstrated an accuracy of 87 % with both optimization methods (Figure 5 (d)). We further analysed the predicted false negative genes and probed their proteomes to investigate the existence of human orthologs (see *Materials and Methods* and Supplementary Table S4). With this, we aimed to eliminate crosslinkings to human-similar proteins since metabolic pathways or enzymes that are missing from the human host have been an important resource of druggable targets against infectious diseases^53^. From the 37 genes that our model predicted to be essential (with FBA and MOMA) contradicting the experimental results, 17 were found to be non-homologous (see Supplementary Table S4). Some examples are the genes encoding the enolpyruvylshikimate phosphate (EPSP) synthase (`A1S_2276`), chorismate synthase (`A1S_1694`), riboflavin synthase (`A1S_0223`), phosphogluconate dehydratase (`A1S_0483`), and 2-keto-3-deoxy-6-phosphogluconate (KDPG) aldolase (`A1S_0484`). The EPSP synthase converts the shikimate3-phosphate together with phosphoenolpyruvate to 5-O-(1carboxyvinyl)-3-phosphoshikimic acid. Subsequently, the chorismate synthase catalyses the conversion of the 5-O-(1carboxyvinyl)-3-phosphoshikimic acid to chorismate, the seventh and last step within the shikimate pathway^54^. Chorismate is the common precursor in the production of the aromatic compounds tryptophan, phenylalanine, and tyrosine, as well as folate and menaquinones during the bacterial life cycle. The shikimate pathway is of particular interest due to its absence from the human host metabolome and its vital role in bacterial metabolism and virulence. Moreover, the enzyme riboflavin synthase catalyses the final step of riboflavin (vitamin B2) biosynthesis with no participating cofactors. Riboflavin can be produced by most microorganisms compared to humans, who have to externally uptake them via food supplements. Also, it plays an important role in the growth of different microbes, especially due to its photosynthesizing property that marks it as a non-invasive and safe therapeutic strategy against bacterial infections^55^. Lastly, the phosphogluconate dehydratase catalyses the dehydration of 6-Phospho-D-gluconate to KDPG, the precursor of pyruvate and 3-Phospho-D-glycerate^56^. This enzyme is part of the Entner–Doudoroff pathway that catabolises glucose to pyruvate, similarly to glycolysis, but using a different set of enzymes^57^.

**Figure 5.**
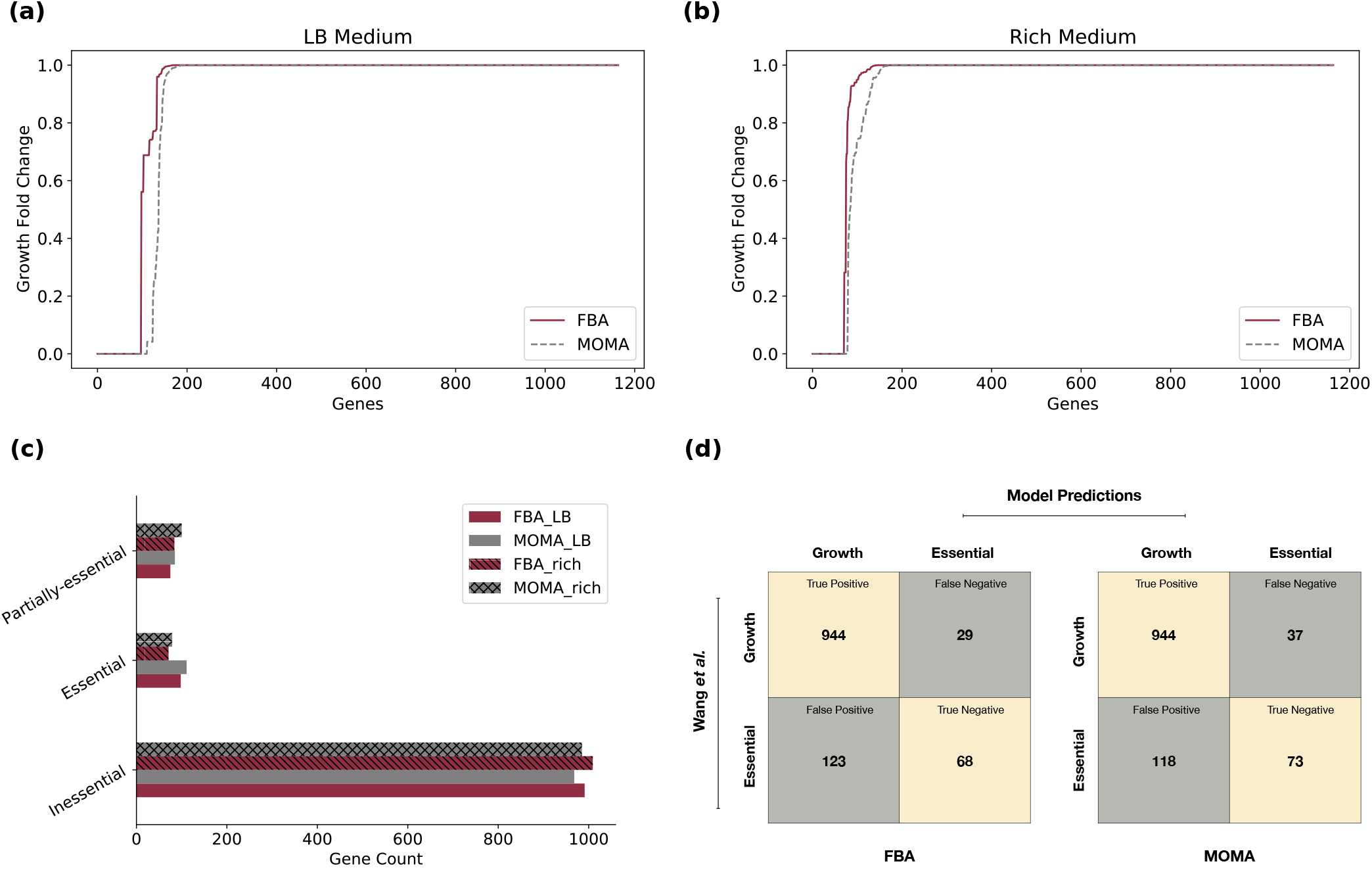
Gene essentiality analysis using *i*ACB23LX. **(a)** and **(b)** Distribution of the *FC*_*gr*_ values calculated for all genes included in *i*ACB23LX. Red lines represent FBA predictions and grey are ratios derived with MOMA. Totally 1,164 knockouts were conducted using each method in the LB medium and rich medium. **(c)** Classification of gene essentialities in LB and the rich growth medium. Both FBA (hatched bars) and MOMA were considered for optimization. The genes were classified as essential, inessential, and partially essential based on their *FC*_*gr*_ values. **(d)** Accuracy of gene essentiality predictions based on empirical data. The *in silico* results were compared to the Wang et al. transponson library. The LB medium was applied to mirror the experimental settings. The metabolic network exhibited 87 % accuracy with FBA (left) and 87 % with MOMA (right). Beige indicates correct predictions; grey indicates incorrect predictions.

We further assessed the druggability of our essential nonhomologous proteins and investigated the existence of inhibitors or compounds known to interact with the enzymes. For this, we used the online DrugBank database that contains detailed information on various drugs and drug targets^58^. In all cases, the listed drugs are of unknown pharmacological action, and there is still no evidence indicating the enzymes’ association with the molecule’s mechanism of action. For instance, the flavin mononucleotide and the cobalt hexamine ion were listed as known inhibitors of yet unknown function against the chorismate synthase, while glyphosate, shikimate3-phosphate, and formic acid have been experimentally found to act with EPSP synthase. Six non-homologous genes were marked as hypothetical or putative in the KEGG^30^ database and/or lacked enzyme-associated information. We searched for drug leads by aligning the query sequences against the DrugBank’s database to find homologous proteins. Two out of six were found to have a protein hit. More specifically, the protein encoded by `A1S_0589` was found to have high sequence identity with the phosphocarrier protein HPr of *Enterococcus faecalis* (Bit-score: 48.5), while the translation product of `A1S_0706` resembles the sugar phosphatase YbiV of *Escherichia coli* (Bit-score: 225.3). According to DrugBank, dexfosfoserine and aspartate beryllium trifluoride have been experimentally determined to bind to these enzymes; however, their pharmacological action is still unknown. The Supplementary Table S4 lists all non-homologous essential genes reported for *i*ACB23LX.

Overall, *i*ACB23LX exhibits high agreement to all validation tests and can, therefore, be used to systematically derive associations between genotypes and phenotypes.

### A curated collection of already published *A. baumannii* metabolic models

In 2010, Kim et al. published the first GEM for the multidrugresistant strain *A. baumannii* AYE^22^. After that, multiple studies provided new data and genomic analyses were published, paving new ways towards its update and refinement^48, 49, 59, 60^. Since then, a variety of GEMs was developed aiming at the empowering of drug development strategies and the enforcement of metabolic engineering by formulating novel and reliable hypotheses (Table 3). However, the amount and format of information contained are inconsistent, with some being syntactically invalid or of older formats. Here, we systematically analysed the quality of all seven currently existing GEMs, reporting their strengths and weaknesses and debugging them to finally build a curated, standardised, and updated collection. To do so, we developed a workflow with curation steps applicable to all models aiming at the standardization and usability of published GEMs by the community (Figure 6 (a)). This closely follows the community-driven workflow published by Carey et al. for the reconstruction of reusable and translatable models^29^. The curation procedure includes a series of stages aiming at modifying data format, data amount, and information quality. It is important to note that no contextual modifications were conducted that could affect the model’s prediction capabilities (see *Materials and Methods*).

**Table 3.** List of genome-scale metabolic models curated for *A. baumannii*, along with information relevant to the manual refinement. Default growth rates (i.e., model simulated as downloaded), the cellular compartments (C: cytosol, E: extracellular space, P: periplasm, and ER: endoplasmic reticulum), and the reactions and metabolites identifiers are listed in the table. MEMOTE scores before and after manual curation are given in the last column. Blue highlights the novel reconstruction for the strain ATCC 17978 presented in this publication. After manual curation, our model developed following our workflow in Figure 1 has the highest quality score and comes along with a minimal medium defined.

|  | Availability | Used Identifiers | Growth by default<br>mmol/(g<sub>DW</sub> · h) | Compartments | MEMOTE |
| --- | --- | --- | --- | --- | --- |
| **AbyMBEL891**<sup>22</sup> | BioModels | Customised | 119 | Cell | 20 % + 17 % |
| **AGORA**<sup>24</sup> | VMH | VMH | 134 | C, E | 42 % + 37 % |
| **iLP844**<sup>23</sup> | Suppl. Mat | ModelSeed | 15.88 | C, E, P | 37 % + 21 % |
| **iCN718**<sup>25</sup> | BiGG | BiGG | 1.31 | C, E, P, ER | 70 % + 3 % |
| **iATCC1906**<sup>26</sup> | Suppl. Mat | KEGG | 46.34 | C, E | 14 % + 44 % |
| **iJS784**<sup>28</sup> | GitHub | ModelSeed | 0.0 | C, E, P | 41 % + 18 % |
| **iAB5075**<sup>27</sup> | Suppl. Mat | BiGG | 1.729 | C, E, P | 17 % + 50 % |
| **iACB23LX** | BioModels | BiGG | 0.1677 | C, E, P | 89 % |

**Figure 6.**
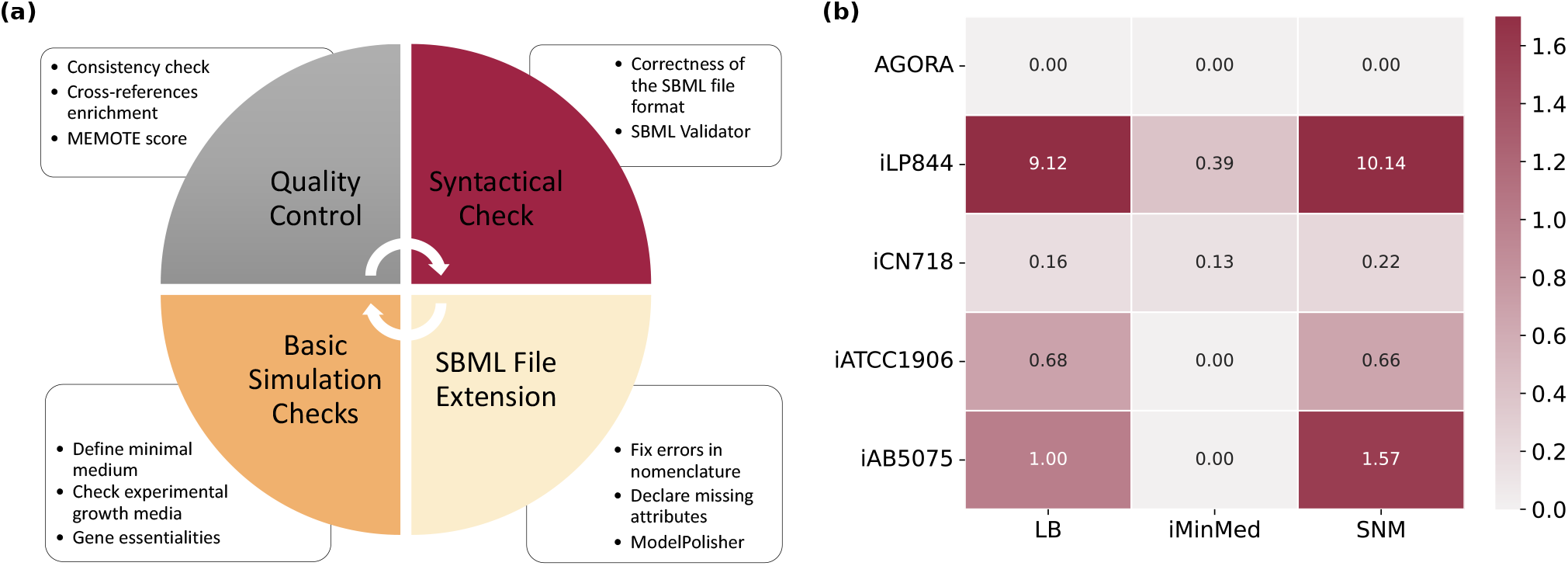
**(a)** Debugging workflow to curate and evaluate already published models. Following the community standards, the already published *A. baumannii* models were curated and transformed into re-usable, simulatable, and translatable models. Quality controls and metabolic standardised tests were conducted using Metabolic Model Testing (MEMOTE), while the validity of the file format and syntax were examined with the SBML Validator^38^. ModelPolisher enhanced the models with missing metadata. **(b)** *In silico*-derived growth rates in various media. The respective growth rates of *i*ACB23LX and the empirical growth rates are shown in Table 2.

Five *A. baumannii* strains have been created throughout the years, with AYE and ATCC 19606 having two reconstructions each. All models are publicly stored and can be downloaded either from a database/repository (BioModels, Virtual Metabolic Human (VMH)^61^, BiGG^62^, and GitHub) or directly from the publication’s supplementary material. The use of distinct identifiers prevents the metabolic networks from being compared to each other. More specifically, *i*LP844 and *i*JS784 carry ModelSEED^63^ identifiers for reactions and metabolites, while *i*CN718 and *i*AB5075 BiGG ^62^ identifiers. AbyMBEL891 uses distinct identifiers not supported by any database, and *i*ATCC1906 includes identifiers derived from KEGG^30^. Most of the models resulted in an unrealistic and inflated growth rate (reference: doubling time of the fastest growing organism *V. natriegens*) in their defined medium, while *i*JS784 showed a zero growth even when all imports were enabled. Hence, this model was excluded from further analysis (Table 3). For each of the remaining GEM, we defined the minimum growth requirements that result in a nonzero and realistic objective value. For instance, the AGORA model required at least 21 compounds (mostly metal ions), while oxygen was sufficient for AbyMBEL891 to simulate a non-zero growth (see Supplementary File S5).

Since these models should successfully reflect the *A. baumannii* metabolic and growth capabilities (see Supplementary Figure S2), we examined the flux through their biomass reaction in various growth media known to induce *A. baumannii* growth (Figure 6 (b)). The majority resulted in a biomass flux of 0.0 mmol/(g_DW_ *·* h) in the *i*MinMed, while the AGORA model could not simulate growth in the LB and SNM as well. Thus, we investigated and identified minimal medium supplementations needed to enable cellular biomass production. As already mentioned, *i*JS784 was excluded from further examination (Table 3), together with AbyMBEL891 that debilitated the analysis due to its non-standardised identifiers and its missing genes and GPRs. When the medium of *i*ATCC1906 and *i*AB5075 was supplemented with D-alanine and D-glucose 6phospahte as well as guanosine 5’-phosphate (GMP), respectively, their biomass reactions carried a positive flux rate of 0.5279 mmol/(g_DW_*·*h) and 0.6477 mmol/(g_DW_ *·*h). Supplementation of meso-2,6-diaminoheptanedioate, menaquinone8, niacinamide, heme, siroheme, and spermidine into the LB medium of the AGORA model resulted in a positive growth rate (1.9430 mmol/(g_DW_ *·*h)). Similarly, when supplementing the SNM with glycyl-L-asparagine, the derived growth rate was 1.5020 mmol/(g_DW_ *·*h), while the *i*MinMed needed to be extended with 12 additional components (resulted growth rate: 1.2789 mmol/(g_DW_*·*h)). Lastly, like with *i*ACB23LX, the LB medium, together with FBA and MOMA, were applied to detect lethal genes in all models (see Supplementary Tables S6 and S7). Despite significant efforts, we could not derive a mapping scheme between the strain-specific gene identifiers of *i*LP844 and *i*ATCC1906 to resolve PROKKA or HMPREF identifiers. Thus, a strain-wise comparison of essential genes would be feasible only for the strain ATCC 17978. As already mentioned, *i*JS784 simulated continuously zero growth and was excluded from the analysis. Consequently, we examined which genes were necessary for growth among the remaining models across three different strains: AYE (*i*CN718), ATCC 17978 (*i*ACB23LX), and AB0057 (AGORA). Totally, 392 genes were identified as essential, while 34 occurred in all three strains. For instance, when the genes encoding for dephospho-coenzyme A (CoA) kinase, phosphopanteth-einyl transferase, shikimate kinase (`A1S_3190`), or chorismate synthase (`A1S_1694`) were deleted from the three strains, no growth could be simulated in the LB medium. As already mentioned, the gene encoding the chorismate synthase has no human-like counterpart. This, together with the fact that it was detected to be vital for growth across three distinct strains, increases its potential to be a drug candidate for future therapies. Generally, most essential genes are members of the purine metabolism and encode various transferases. Besides this, the pantothenate and CoA biosynthesis and the amino acid metabolism were found to be a prominent target pathways for further drug development.

## Discussion

The historical timeline of past pandemics shows the imposed threat of bacteria in causing repetitive outbreaks with the highest death tolls^64^, such as cholera and plague. By 2050, antimicrobial-resistant pathogens are expected to kill 10 million people annually^65^, while the antibiotics misuse accompanied by the ongoing Coronavirus Disease 2019 (COVID-19) crisis exacerbated this global threat. It is noteworthy that elevated morbidity rates were ascribed to bacterial co/secondary infections during previous viral disease outbreaks^66, 67, 68^. Hence, developing effective antibiotic regimens is of urgent importance. Here we present the most recent and comprehensive ready-to-use blueprint GEM for the Gram-negative pathogen *A. baumannii*. For this, we developed a workflow that applies to any living organism and ensures the reconstruction of high-quality models following the community standards. Our model, *i*ACB23LX, was able to simulate growth in the SNM that mimics the human nasal niche, the experimentally defined medium LB, and the model-derived *i*MinMed. With *i*MinMed we denoted the minimal number of compounds needed to achieve non-zero growth. This medium contains totally 14 compounds, including transition metals and energy sources. Transitions metals have been shown to participate in important biological processes and are vital for the survival of living organisms^41^. We confirmed the computationally predicted growth rates by comparing them to our empirically determined growth kinetics data. With this, we ensured that our model recapitulates growth phenotypes in media that reflect *Acinetobacter*-associated environments.

Furthermore, we validated *i*ACB23LX quantitatively and qualitatively using existing experimental data and observed remarkable improvements compared to precursory models. More specifically, our model predicted experimental Biolog growth phenotypes on various carbon sources^48^ with 86.3 % overall agreement, which is higher than the predictions capability of *i*ATCC19606 (84.3 %) and *i*LP844 (84 %), and comparable to that of *i*AB5075 (86.3 %). Similarly, *i*ACB23LX exhibited 79.2 % predictive accuracy on nitrogen sources tests, while this increases to 87.5 % after further refinement.Improving and re-defining the biomass objective function (BOF) based on accurate strain-specific experimental data would be the next step to diminish the number of inconsistent predictions and to further improve the network and its predictive potential. During gene lethality analysis in LB medium, our model predicted 110 genes with MOMA to be essential, while 97 of them were also reported by FBA to impair the growth. Generally, after enriching the nutritional input with all available compounds (rich medium), less lethal genes resulted, meaning that *A. baumannii* undergoes metabolic alterations when nutrients are lacking. Our *in silico* results compared to the strain-specific gene essentiality data^49^ resulted in 87 % overall accuracy, which is remarkably higher than all GEMs built for *A. baumannii* (e.g., 80.22 % for *i*CN718 and 72 % for *i*LP844), except *i*AB5075 which performed comparably. Subsequently, we examined more carefully our false negative predictions and searched for putative drug targets that could be employed for future therapeutics. More specifically, we focused on genes found to be essential for growth and encode proteins with no human counterparts (see Supplementary Table S4). Our study highlighted the EPSP and chorismate synthases from the shikimate pathway as prominent target candidates with no correlation to the human proteome. Several knockout studies have highlighted the significance of enzymes from the shikimate metabolism as potential targets against infections caused by threatening microorganisms, e.g., *Mycobacterium tuberculosis*^69^, *Plasmodium falciparum*^70^, and *Yersinia enterocolitica*^71^. Umland et al. identified these two gene products as essential in an *in vivo* study using a clinical isolate of *A. baumannii* and a rat abscess infection model^72^. This increases the confidence of our results and indicates that novel genes found to be essential *in silico* should be considered as potential antimicrobial targets. Similarly, numerous studies have suggested one of our further candidates, riboflavin, as a potential antimicrobial agent^55^, while the Entner–Doudoroff pathway (in which our candidate targets phosphogluconate dehydratase and KDPG aldolase act to produce pyruvate) is similar to the glycolysis but with different member enzymes, has been firstly discovered in *Pseudomonas saccharophila*^57^ and later in *E. coli*^73^. Meanwhile, it is vital for the survival of further pathogenic microorganisms, like *Neisseria gonorrhoeae, Klebsiella pneumoniae*, and *Pseudomonas aeruginosa*^74, 75, 76^. However, they have not yet been examined in the context of *Acinetobacter* species and could be a source of antimicrobial therapeutic strategies. Hence, these biosynthetic routes could be a valuable resource for targets to fight bacterial infectious diseases. Finally, we investigated the druggability of our essential nonhomologous genes. We searched the DrugBank database to find compounds known to inhibit these genes and that are already approved by the Food and Drug Administration (FDA). Our analysis resulted in only drugs that have been found to interact with the gene product of interest; however their pharmacological action is yet unknown. We further probed the hypothetical and putative non-homologous genes against the DrugBank’s sequence database to find homologous proteins and determine their activity. Also in this case, the resulted drugs were listed with still undetermined pharmacological action. These putative and yet unexplored targets with inhibitory potential are of great interest in the context of developing novel classes of antibiotics. Overall, our model reached a `MEMOTE` score of 89 %, which is the highest score reported for this organism.

Moreover, we improved and assessed all previously published models and created the first curated collection of metabolic networks for *A. baumannii*. We created a debugging workflow consisting of four major steps to systematically analyse and curate constraint-based models focusing on their standardization and the FAIR data principles. We applied this workflow and curated a total of seven metabolic models for *A. baumannii*. In addition, most of the models simulated growth rates by default that were unrealistic when compared to the fastest growing organism (*V. natriegens*)^36^. Hence, we determined the minimal number of components needed for these models to result in non-inflated biomass production rates. The defined minimal media were mostly composed of metal ions (e.g., cobalt, iron, magnesium) that are essential for bacterial growth. For the model *i*JS784, the minimization process was infeasible; thus, the model was not considered for further analysis. We also examined the growth ability of these models in three media (SNM, LB, and *i*MinMed) and compared them to our model, *i*ACB23LX. When the models simulated a zero flux through the biomass reaction, we continued by detecting the minimal amount of metabolites supplemented in the medium that resulted in a non-zero growth rate. These would enable the detection of gaps and assist in future improvement of the models. It is important to note here that with this curation, we opted for a systematical assessment of the previously reconstructed models and the detection of their assets and liabilities. Consequently, we did not undertake any contextual modification that could alter the models’ predictive capabilities. Finally, we *in silico* detected lethal genes among comparable and simulatable models of *A. baumannii*. Our analysis incorporated three strains of *A. baumannii* (AYE, AB0057, and ATCC 17978), and we examined the effect of genetic variation across strains in the gene essentiality. Our analysis highlighted once again the shikimate pathway, as well as the purine metabolism, the pantothenate, and CoA biosynthesis, and the amino acid metabolism as candidate routes to consider for future new classes of antibacterial drugs with potential effect across multiple *A. baumannii* strains. The curated models, together with our novel model, would benefit the future prediction of candidate lethal genes by reducing the considerable resources needed for classical whole-genome essentiality screenings. All in all, this collection of simulation-ready models will forward the selection of a suitable metabolic network based on individual research questions and help define the entire species and new hypothesis.

Our new metabolic reconstruction and the curated collection of further strain-specific models will guide the formulation of ground-breaking and reliable model-driven hypotheses about this pathogen and help examine the diversity in the metabolic behavior of different *A. baumannii* species in response to genetic and environmental alterations. Additionally, they can be utilised as knowledge bases to detect critical pathways related to responses against multiple antibiotic treatments. This will ultimately strengthen the development of advanced precision antimicrobial control strategies against multidrugresistant (MDR) *A. baumannii* strains.

Taken together, our workflows and models can be employed to expand this collection further with additional standardised strain-specific metabolic reconstructions to finally define the core and pan metabolic capabilities of *A. baumannii*.

## Materials and Methods

### Growth curves of *A. baumannii*

Growth curves for *A. baumannii* strains AB5075, ATCC 17978, ATCC 19606 and AYE were recorded in LB medium, *i*MinMed medium supplemented with acetate as the sole carbon source (0.2 % weight per volume, respectively), and SNM^77^. Overnight cultures of the strains grown in LB medium were harvested by centrifugation, and the cell pellet was washed once with 5 mL of phosphate buffered saline (PBS). Cells were then re-suspended in the medium used for the growth curves. The starting optical density (OD)_600 nm_ was adjusted to 0.1 and growth curves were recorded in 2 mL of medium for 12 h in triplicates using a Tecan infinite M200 PRO plate reader and 12-well plates covered with a plastic lid. Plates were incubated with linear shaking at 37 ^*°*^C and the OD_600 nm_ was measured every 15 min. Growth rates were determined as the slope of the linear part of the curves plotting the natural logarithm of OD_600 nm_ against time.

### The metabolic model reconstruction workflow

Figure 1 illustrates the workflow we developed to create the novel high-quality genome-scale metabolic network *i*ACB23LX, following the state-of-the-art protocol of Thiele and Palsson^20^. Our workflow consists of eight major steps starting from the extraction of an annotated genome until the model validation using experimental data. Modifications in the model structure, as well as the inclusion of cross-references to multiple functional databases, were done using the libSBML^38^ library, while all simulations were conducted via the COBRApy -0.22.1^50^ suite that includes functions commonly used for simulations.

The individual steps are described below in more detail with respect to the reconstruction of *i*ACB23LX.

### Draft reconstruction

A first draft model was built with CarveMe version 1.5.1 using the annotated genome sequence of the strain ATCC 17978. This was downloaded from the National Centre for Biotechnology Information (NCBI) at https://www.ncbi.nlm.nih.gov and has the assembly accession number `ASM1542v1`^78^. Seven strain-specific assemblies are registered in NCBI; however, the chosen entry is also present in the KEGG^30^ database facilitating the model extension. The genome is 3.9 Mbp long and has two plasmids (pAB1 and pAB2). We set the SBML flavor to activate the extension for fbc version 2^79^ that allows semantic descriptions for domain-specific elements such as metabolite chemical formulas and charges together with reaction boundaries and GPRs. Moreover, the CarveMe parameter `gramneg` was selected to employ the specialised template for the Gram-negative bacteria. Compared to the Gram-positive template, the Gram-negative template comes with phosphatidylethanolamines, murein, and a lipopolysaccharide unit. Its biomass reactions involve membrane and cell wall components resulting in more accurate gene essentiality predictions in the lipid biosynthesis pathways.

### Manual refinement and extension

We started the manual refinement of the draft model by resolving syntactical errors within the model file using SBML Validator from the libSBML library^38^. Missing metabolite charges and chemical formulas were retrieved from the BiGG^62^ and ChEBI^80^ databases, while massand chargeimbalanced reactions were corrected. The most intense part of the workflow is the manual network extension and gapfilling. This was done using the organism-specific databases KEGG^30^ and BioCyc^81^, together with ModelSEED^63^. We mapped the new gene locus tags to the old ones using the GenBank General Feature Format (GFF)^82^ and added missing metabolic genes along with the respective reactions and metabolites into our model. The network’s connectivity was ensured by resolving as many dead-ends (are only produced but not consumed) and orphan (are only consumed but not produced) metabolites as possible. Also, reactions with no connectivity were not included in the model, while reactions with no organism-specific gene evidence were removed from the model.

### Erroneous energy generating cycles

Energy generating cycles (EGC) are thermodynamically infeasible loops found in metabolic networks and have not been experimentally observed, unlike futile cycles. EGCs charge energy metabolites like adenosine triphosphate (ATP) and uridine triphosphate (UTP) without any external source of nutrients and may result in incorrect and unrealistic energy increases. Their elimination is crucial while correcting the energy metabolism since they can inflate the maximal biomass yields and make the predictions unreliable. We checked their existence in *i*ACB23LX applying an algorithm developed by Fritzemeier et al.^32^.

We created a Python script that (1) defines and adds energy dissipation reactions (EDRs) in the network:

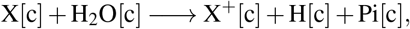

where X is the metabolite of interest and (2) maximises each EDR while blocking all influxes. This can be formulated as follows:

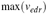

subject to

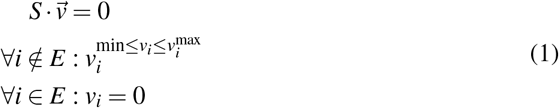

where *edr* is the index of the current dissipation reaction, *S* is the stoichiometric matrix, 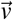 the flux vector, *E* the set of all exchange reactions, and 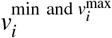 the upper and lower bounds. The existence of EGCs is indicated by a positive optimal value of *v*_*edr*_.

Totally we examined 14 energy metabolites: ATP, cytidine triphosphate (CTP), guanosine triphosphate (GTP), UTP, inosine triphosphate (ITP), nicotinamide adenine dinucleotide (NADH), nicotinamide adenine dinucleotide phosphate (NADPH), flavin adenine dinucleotide (FADH2), flavin mononucleotide (FMNH2), ubiquinol-8, menaquinol-8, demethylmenaquinol-8, acetyl-CoA, and L-glutamate. Moreover, we tested the proton exchange between cytosol and periplasm.

In the case of existing EGCs, we examined the directionality and the gene evidence of all participated reactions using the BioCyc organism-specific information as reference^81^.

### Database annotations

In this stage, the model was enriched with cross-linkings to various functional databases. Reactions and metabolites were annotated with databases (e.g., KEGG^30^, BRENDA^83^, and UniProt^84^). These were included in the model as controlled vocabulary (CV) Terms following the Minimal Information Required In the Annotation of Models (MIRIAM) guidelines^85^ and the resolution service at https://identifiers.org/. We used ModelPolisher^86^ to complete the missing available metadata for all metabolites and genes. Similarly, metabolic genes were annotated with their KEGG^30^, NCBI Protein, and RefSeq identifiers using the GFF^82^. To reactions, metabolites, and genes SBO terms were assigned using the SBOannotator^34^. SBO terms provide unambiguous semantic information and specify the type or role of the individual model component^87^. In addition, ECO terms were added to every reaction to capture the type of evidence of biological assertions with `BQB_IS_DESCRIBED_BY` as a biological qualifier. They are useful during quality control and mirror the curator’s confidence about the inclusion of a reaction. When multiple genes encode a single reaction, an ECO term was added for every participant gene. Both terms were incorporated into the model according to our mapping in Figure 3.

Finally, reactions were annotated with the associated subsystems in which they participate using the KEGG^30^ database and the biological qualifier `BQB_OCCURS_IN`. Moreover, the “groups” plugin was activated^39^. Every reaction that appeared in a given pathway was added as a `groups:member`, while each pathway was created as a group instance with `sboTerm=“SBO:0000633”` and `groups:kind=“partonomy”`.

### Quality control and quality assurance

`MEMOTE`^36^ version 0.13.0 was used to assess and track the quality of our model after each modification, providing us with information regarding the model improvement. The final model was converted into the latest SBML Level 3 Version 2^39^ format using the libSBML package, while the SBML Validator tracked syntactical errors and ensured a valid format of the final model^38^.

### Constraint-based analysis

The most frequently used constraint-based modelling approach is the FBA that determines a flux distribution via optimization of the objective function and linear programming^51^. Prior to this, the metabolic network is mathematically encoded using the stoichiometric matrix *S* formalism. This structure delineates the connectivity of the network, and it is formed by the stoichiometric coefficients of all participating biochemical reactions. The rows and columns are represented by the metabolites and the massand charge-balanced reactions respectively. At steady state, the system of linear equations derived from the network is defined as follows:

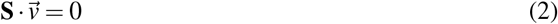

with **S** being the stoichiometric matrix and 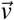 the flux vector. With no defined constraints, the flux distribution may be determined at any point within the solution space. This space must be further restricted since the system is under-determined and algebraically insoluble. An allowable solution space is defined by a series of imposed constraints that are followed by cellular functions. Altogether the FBA maximization problem, with mass balance, thermodynamic, and capacity constraints, is defined as:

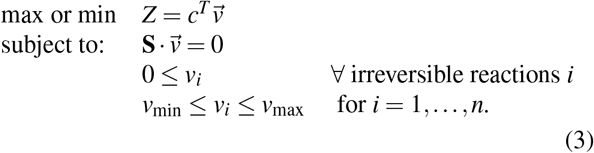

Here, *n* is the amount of reactions, *Z* represents the linear objective function, and 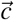 is a vector of coefficients on the fluxes 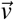 used to define the objective function.

### Growth simulations

#### Strict aerobic growth check

At the time of writing, the utilised draft reconstruction tool, CarveMe^31^, does not include reconstruction templates to differentiate between aerobic and anaerobic species. The directionality of reactions that produce or consume oxygen may affect the model’s ability to grow anaerobically. *A. baumannii* is defined to be a strictly aerobic species. Hence, we tested whether our model could grow with no oxygen supplementation. For this purpose, we examined all active oxygen-producing reactions under anaerobic conditions. We corrected their directionality based on the organism-specific information found in BioCyc^81^ and kept only those with associated gene evidence.

#### Defining a minimal growth medium

To determine the minimal number of nutrients needed for the bacterium to grow, we defined a minimal medium using *i*ACB23LX. We determined the minimal amount of metabolites needed for growth using the M9 medium (Supplementary File S1) as a reference. We modeled growth on the minimal medium by enabling the uptake of all metabolites that constitute the medium (lower bound of exchange reactions set to −10 mmol/(g_DW_ *·* h)). The lower bound for the rest of the exchanges was set to 0 mmol/(g_DW_ *·* h). The final minimal medium is listed in Table 1 and the Supplementary File S1. It consists of nine transition metals, a carbon source, a nitrogen source, a sulfur source, and a phosphorus source. The aerobic environment was simulated by setting the lower bound for the oxygen exchange to −10 mmol/(g_DW_ *·* h).

#### Growth in chemically defined media

We utilised experimentally verified growth media to examine the growth capabilities of *i*ACB23LX. The LB medium serves as a common medium for the cultivation of *A. baumannii*. Consequently, we conducted an assessment of our model’s capacity to accurately simulate growth in this particular medium Additionally, we inspected the growth of our model in the human nasal niche, as *A. baumannii* have been isolated from nasal samples within ICUs^10, 11, 12^. For this purpose, we utilised the SNM that imitates the human nasal habitat^77^. In all cases, if macromolecules or mixtures were present, we considered the constitutive molecular components for the medium definition. As our model was initially unable to reproduce growth on the applied media, we deployed the gap-filling option from CarveMe to detect missing reactions and gaps in the network^31^. All growth media formulations are available in the Supplementary File S1.

#### Rich medium definition

To investigate our model’s growth rate when all nutrients are available to the bacterial cell, we defined the rich medium. For this purpose, we enabled the uptake of all extracellular metabolites by the model setting the lower bound of their exchange reactions to −10 mmol/(g_DW_ *·* h).

### Model validation

#### Evaluation of carbon and nitrogen utilization

We employed the previously published Biolog Phenotypic Array data by Farrugia et al. for *A. baumannii* ATCC 17978 to validate the functionality of our model^48^. According to the experimental guidelines provided by Farrugia et al., we utilised the M9 medium for all simulations. The medium was then supplemented with D-xylose as a carbon source for the nitrogen testings, while ammonium served as the only nitrogen source for the carbon tests. As D-xylose was initially not part of the model, we conducted an extensive search in the organism-specific databases KEGG^30^ and BioCyc^81^ to include missing reactions.

The phenotypes were grouped by their maximal kinetic curve height. A trait was considered positive (“growth”) if the height exceeded the 115 and 101 OmniLog units for a nitrogen and carbon source, respectively. The prediction accuracy was evaluated by comparing the *in silico*-derived phenotypes to the Biolog results. More specifically, the overall model’s accuracy (ACC) was calculated by the overall agreement:

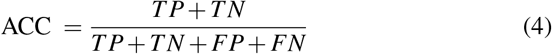

where true positive (TP) and true negative (TN) are correct predictions, while false positive (FP) and false negative (FN) are inconsistent predictions. Discrepancies were resolved via iterative manual curation of the model.

#### Gene perturbation analysis

We performed *in silico* single-gene deletions on *i*ACB23LX to detect essential genes. For this purpose, we utilised the `single_gene_deletion` function from the COBRApy^50^ package. A gene is considered to be essential if a flux of 0.0 mmol/(g_DW_ *·*h) was observed through the biomass reaction after setting the lower and upper bounds of the associated reaction(s) to 0.0 mmol/(g_DW_ *·*h).

Additionally, we examined the effect of gene deletions using two different optimization approaches: FBA^51^ and MOMA^52^. Contrary to FBA, MOMA is based on quadratic programming, and the involved optimization problem is the Euclidean distance minimization in flux space. Moreover, it approximates the metabolic phenotype and relaxes the assumption of optimal growth flux for gene deletions^52^.

The results were compared to the ATCC 17978-specific gene essentiality dataset from 2014^49^. Wang et al. generated a random mutagenesis dataset including 15,000 unique transposon mutants using insertion sequencing (INSeq)^49^. By the time of writing, this is the only *A. baumannii* ATCC 17978 library presenting gene essentiality information. Analogously to the experimental settings, the nutrient uptake constraints were set to the LB medium. From the 453 genes identified as essential by Wang et al., 191 could be compared to our predictions. The rest were not part of *i*ACB23LX due to their non-metabolic functions. To measure the effect of a single deletion, we calculated the fold change (FC) between the model’s growth rate after (*gr*_*KO*_) and before (*gr*_*WT*_) a single knockout. This is formulated as follows:

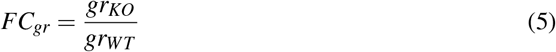

To this end, if *FC*_*gr*_ = 0, the deleted gene is classified as essential, meaning its removal prevented the network from producing at least one key biomass metabolite predicting no growth. Similarly, if *FC*_*gr*_ = 1, the deletion of the gene from the network did not affect the growth phenotype (labeled as inessential), while when 0 < *FC*_*gr*_ < 1, the removal of this gene affected partially the biomass production (labeled as partially essential). The complete lists of the gene essentiality results are available in the Supplementary Files S2 and S3.

To examine the potential of the *in silico* determined essential genes on becoming novel drug candidates to fight *A. baumannii* infections, we probed the queries of predicted false negative candidates against the human proteome using Basic Local Alignment Search Tool (BLAST)^88^. The protein sequences were aligned to the human protein sequences using the default settings of the NCBI BLASTp tool (word size: 6, matrix: BLOSUM62, gap costs: 11 for existence and 1 for extension). To eliminate adverse effects and ensure no interference with human-like proteins, queries with any non-zero alignment score with the human proteome were not considered. Lastly, we searched in the DrugBank database version 5.1.9 to find inhibitors or ligands known to act with the enzymes encoded by the non-homologous genes^58^.

#### Curation of existing metabolic networks

Previously reconstructed models of *A. baumannii* for multiple strains (Figure 2, Table 3) were collected and curated following community standards and guidelines. For this, we created a workflow shown in Figure 6 that consists of four main steps and utilises model validation and annotation tools. This can be applied to any metabolic network in SBML^37^ format and follows the community “gold standards” strictly as proposed by Carey et al.^29^. The curation steps involved changes in the format, amount, and quality of the included information. The context has not been altered in any way that could impact the models’ prediction capabilities. We employed a combination of already existing tools to analyse, simulate, and quality-control the models (COBRApy^50^, `MEMOTE`^36^, and the SBML Validator^38^). Different database cross-references were incorporated in the models using ModelPolisher^86^ and following the MIRIAM guidelines^85^, while the libSBML library^38^ was used to manipulate the file format and convert to the latest version. To resolve inflated growth rates, we determined computationally-defined minimal growth media. The growth capabilities were examined with respect to various experimentally-derived growth media, while the LB medium was applied to identify lethal genes. A strain-wise comparison was not feasible due to strain-specific identifiers, no successful growth, or missing genes. Hence, we investigated the essential genes across all models with identifiers that could not be mapped with the Pathosystems Resource Integration Center (PATRIC) ID mapping tool^89^.

To begin with the debugging, we examined the syntactical correctness and internal consistency of the downloaded files using the SBML Validator from the libSBML library^38^. Two models (*i*CN718 and *i*JS784) could not pass the validator check and reported errors since they were not in a valid SBML^37^ format right after their attainment. We made *i*CN718 valid by deleting the reaction `DNADRAIN` for which neither a reactant nor a product was assigned since the associated metabolite was not part of the model. Similarly, the empty `groups` attribute was removed from *i*JS784, converting the file into a valid format. Warnings were detected for *i*ATCC1906, and *i*AB5075 due to missing definition of the fbc extension (became available at the latest Level 3 release^79^) and the non-alphanumeric chemical formulas. We resolved these issues by defining the fbc list `listOfGeneProducts` and the species attribute `chemicalFormula`. In more detail, we extracted the given GPR from the notes field and defined individual `geneProduct` classes with `id`, `name`, and `label`. The attribute `chemicalFormula` was set equal to the species chemical formulas extracted from the notes and is particularly essential in reaction’s validation and balancing. Following the SBML^37^ specifications regarding its constitution, in case of ambiguous formulas separated by a semicolon (;), the first molecular representation was chosen. With this, the genes and metabolites’ chemical formulas became part of the file’s main structure. Since *i*ATCC1906 carried KEGG^30^ identifiers, we could extract the metabolites’ chemical formulas from the database and add them to the model. Moving on with the file extension, we declared the remaining missing attributes from reactions, metabolites, and genes that are required according to the SBML^37^ language guidelines. More specifically, we defined the `metaid` attribute when missing, while we fixed any errors regarding the identifiers nomenclature. Further extension involved the annotation of reactions, metabolites, and genes with a plethora of database cross-references following the MIRIAM guidelines^85^. For this, we employed ModelPolisher that complements and annotates SBML^37^ models with additional metadata using the BiGG Models knowledgebase as reference^86^. We also defined precise SBO Terms with the `sboTerm` attribute using the SBOannotator^34^. The final step of debugging involved the conversion of all models to the newest available format SBML Level 3 Version 2^39^, as well as the quality control using `MEMOTE`^36^.

## Data availability

Supplementary tables in Microsoft Excel format are available along with this article. The model *i*ACB23LX along with all curated and refined models are available at the BioModels Database^90^ as an SBML Level 3 Version 1^91^ file distributed as Open Modelling EXchange format (OMEX) archive^92^ including annotation^93^. Access the model at https://www.ebi.ac.uk/biomodels/MODEL2309120001.

## Author contributions

Conceptualization and idea, N.L.; model reconstruction and analysis, N.L. and Y.X.; curation and analysis of additional models, N.L.; experimental work, L.F. and M.S.; manuscript writing, N.L.; manuscript revision, N.L., Y.X., and A.D.; supervision and funding acquisition, A.D. All authors approved the publishing of the manuscript.

## Acknowledgments

This work was funded by the *Deutsche Forschungsgemeinschaft* (DFG, German Research Foundation) under Germany’s Excellence Strategy – EXC 2124 – 390838134 and supported by the Cluster of Excellence ‘Controlling Microbes to Fight Infections’ (CMFI). A.D. is supported by the German Center for Infection Research (DZIF, doi: 10.13039/100009139) within the *Deutsche Zentren der Gesundheitsforschung* (BMBF-DZG, German Centers for Health Research of the Federal Ministry of Education and Research (BMBF)), grant 𝒩0 8020708703. The authors acknowledge the support by the Open Access Publishing Fund of the University of Tübingen (https://uni-tuebingen.de/en/216529). The authors also thank Dr. Bernhard Krismer for providing the synthetic nasal medium.

## List of Abbreviations

AGORA: Assembly of Gut Organisms through Reconstruction and Analysis
AMR: antimicrobial resistance
ATP: adenosine triphosphate
BiGG: Biochemical, Genetical, and Genomical
BLAST: Basic Local Alignment Search Tool
BMBF: Federal Ministry of Education and Research (*Bundesministerium für Bildung und Forschung*)
BMBF-DZG: *Deutsche Zentren der Gesundheitsforschung*
BOF: biomass objective function
CBM: constraint-based modelling
CMFI: Controlling Microbes to Fight Infections
COBRApy: Constraints-Based Reconstruction and Analysis for Python
CoA: coenzyme A
COVID-19: Coronavirus Disease 2019
CTP: cytidine triphosphate
CV: controlled vocabulary
DFG: *Deutsche Forschungsgemeinschaft*
DZIF: German Center for Infection Research
ECO: Evidence and Conclusion Ontology
EDR: energy dissipation reaction
EGC: energy-generating cycle
EPSP: enolpyruvylshikimate phosphate
FADH2: flavin adenine dinucleotide
FAIR: Findable, Accessible, Interoperable, and Reusable
FC: fold change
FMNH2: flavin mononucleotide
FBA: flux balance analysis
fbc: flux balance constraints
FDA: Food and Drug Administration
FN: false negative
FP: false positive
GEM: genome-scale metabolic model
GFF: General Feature Format
GMP: guanosine 5’-phosphate
GPR: gene-protein-reaction associations
GTP: guanosine triphosphate
ICU: intensive care unit
INSeq: insertion sequencing
ITP: inosine triphosphate
JSON: JavaScript Object Notation
KDPG: 2-keto-3-deoxy-6-phosphogluconate
KEGG: Kyoto Encyclopedia of Genes and Genomes
LB: Luria-Bertani
M9: M9 minimal medium
MDR: multidrug-resistant
MEMOTE: Metabolic Model Testing
MIRIAM: Minimal Information Required In the Annotation of Models
MOMA: Minimization of Metabolic Adjustment
NADH: nicotinamide adenine dinucleotide
NADPH: nicotinamide adenine dinucleotide phosphate
NCBI: National Centre for Biotechnology Information
OD: optical density
OLS: Ontology Lookup Service
OMEX: Open Modelling EXchange format
PATRIC: Pathosystems Resource Integration Center
PBS: phosphate buffered saline
DR: pandrug-resistant
SARS-CoV-2: Severe Acute Respiratory Syndrome Coronavirus 2
SBML: Systems Biology Markup Language
SBO: Systems Biology Ontology
SNM: synthetic nasal medium
TN: true negative
TP: true positive
UTP: uridine triphosphate
VMH: Virtual Metabolic Human
WHO: World Health Organization
XDR: extensively drug-resistant

## Supporting Information

**S1 Figure. Oxygen-producing and -consuming reactions found in *i*ACB23LX together with their anaerobic fluxes**. All flux rates are written in orange and are given in mmol/(g_DW_ *·*h). The reaction abbreviations are as follows: `O2tpp`, O_2_ transport via diffusion between periplasm and cytosol; `CATpp`, periplasmatic catalase; `H2O2tex`, hydrogen peroxide transport via diffusion; `CAT`, Catalase; `O2tex`, O_2_ transport via diffusion between periplasm and extracellular space; `EX_h2o2_e`, hydrogen peroxide exchange and `EX_o2_e`, O_2_ exchange. Figure generated with Escher^94^.

**S2 Figure. Experimentally-derived growth curves of**

***A. baumannii***. The growth curves for *A. baumannii* strains AB5075, ATCC 73217978, ATCC 19606, and AYE were measured in LB medium, M9 medium supplemented with acetate, and acSNM. Additionally, the *in silico*-defined minimal medium (*i*MinMed) was tested for all strains.

**S1 Table. *In silico* formulations of examined media compositions**. Metabolites are described by BiGG ^62^ identifiers.

**S2 Table. *In silico* gene knockout results using FBA**. The ratio column describes the growth rate change before and after the respective knockout.

**S3 Table. *In silico* gene knockout results using MOMA**. The ratio column describes the growth rate change before and after the respective knockout.

**S4 Table. Metabolic genes found to be essential for growth in *i*ACB23LX and encode proteins with no human counterparts**.

**S5 Table. Computationally-defined minimal growth media for previously published models**. Due to inflated growth rates in most published *A. baumannii* GEMs, we established minimal media supporting non-zero biomass flux.

**S6 Table. Gene lethality predictions using previously published *A. baumannii* models and FBA**.

**S7 Table. Gene lethality predictions using previously published *A. baumannii* models and MOMA**. This offers a complementary perspective on the essential genes in the organism’s metabolism.

