## Supplementary figures and images for "Exploring the metabolic profiling of *A. baumannii* for antimicrobial development using genome-scale modelling"

### S1_Figure.pdf

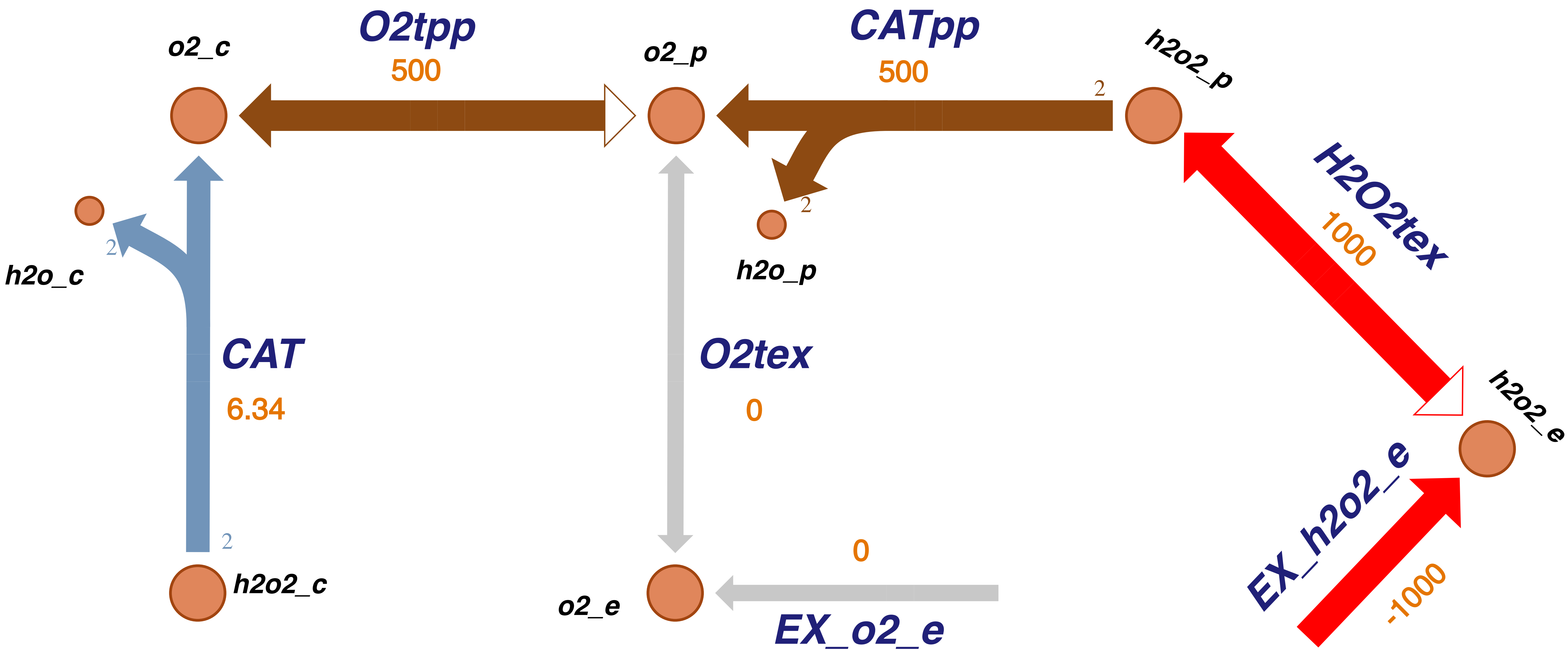

### S2_Figure.pdf

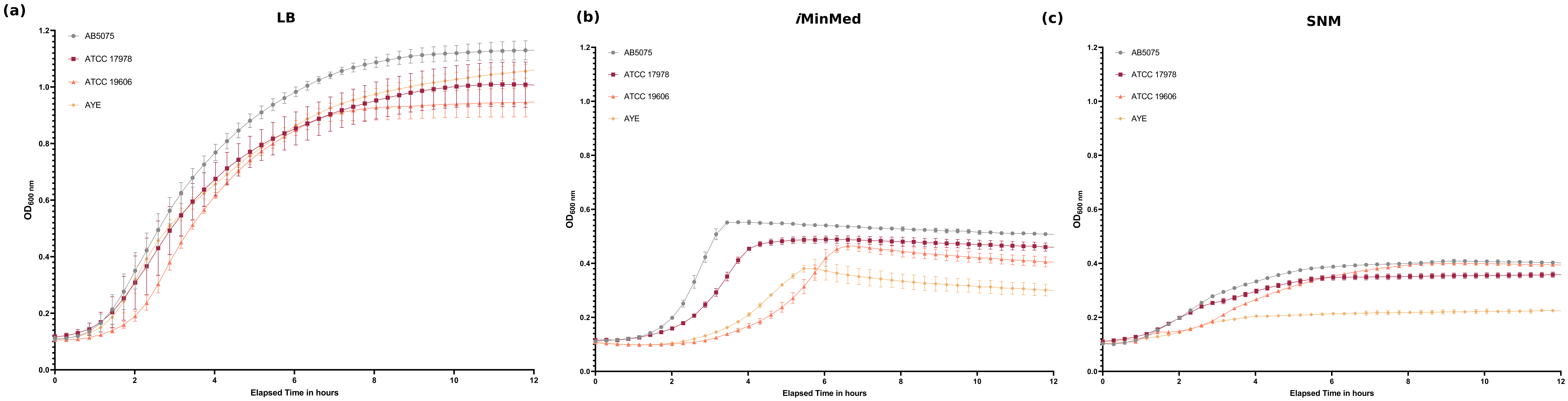
